# YAP and collagen remodelling support cell proliferation and tumour aggressiveness in uterine leiomyosarcoma

**DOI:** 10.1101/2022.06.27.497746

**Authors:** Jordi Gonzalez-Molina, Paula Hahn, Raul Maia Falcão, Okan Gultekin, Georgia Kokaraki, Valentina Zanfagnin, Tirzah Braz Petta, Kaisa Lehti, Joseph W. Carlson

## Abstract

Fibrillar collagen deposition, stiffness, and downstream signalling support the development of leiomyomas (LM), common benign mesenchymal tumours of the uterus, and are associated with aggressiveness in multiple carcinomas. Compared to epithelial carcinomas, however, the impact of fibrillar collagens on malignant mesenchymal tumours, including uterine leiomyosarcoma (LMS), remains elusive. In this study, we analyse the network morphology and density of fibrillar collagens combined with the gene expression of LMS, LM and normal myometrium (MM). We find that, in contrast to LM, LMS tumours present low collagen density and increased expression of collagen-remodelling genes, features associated with tumour aggressiveness. Using collagen-based 3D matrices, we show that the activity of MMP14, a central protein with collagen-remodelling functions particularly overexpressed in LMS, is necessary for LMS cell proliferation. In addition, we find that, unlike MM and LM cells, LMS proliferation and migration are not affected by collagen substrate stiffness. We demonstrate that LMS cell growth in low matrix adhesion microenvironments is supported by an enhanced basal YAP activity. Altogether, our results indicate that LMS cells acquire high collagen remodelling capabilities and are adapted to grow and migrate in low collagen and soft microenvironments. These results further suggest that matrix remodelling and YAP are potential therapeutic targets for this deadly disease.

## Introduction

Leiomyosarcomas (LMS) are malignant mesenchymal neoplasms showing features of smooth muscle lineage that represent 10-20% of all newly diagnosed soft tissue sarcomas (STSs) ^1^. Uterine LMS are the most common subtype of uterine sarcoma. These tumours are generally treated surgically. Despite apparent complete resection of LMS tumours, the risk of recurrence is between 50-70% ^2^. The symptoms of uterine LMS are similar to those of uterine leiomyomas (LM), common benign fibrotic tumours of the uterus, and thus, most LMS are diagnosed postoperatively ^3,4^. Treatment of uterine LM includes tissue morcellation, which can lead to the spreading of LMS in case of misdiagnosis ^5,6^. Understanding the biological basis of LMS development and progression and its differences with LM is fundamental to improve the diagnosis and treatment of this deadly disease.

The desmoplastic reaction is a characteristic increase in extracellular matrix (ECM) deposition and crosslinking, particularly of fibrillar collagens, that occurs in various types of carcinoma ^7^. Similarly, LM are characterised by an enrichment in fibrillar collagens type I and III ^8,9^. The increased deposition, crosslinking, and re-organisation of fibrillar collagens causes the stiffening of the ECM and the bulk tissue ^10^. Increased tissue stiffness is a characteristic of desmoplastic tumours. Likewise, LM tumours present a higher average stiffness (18.6 kPa) than normal myometrium (MM; 4.9 kPa) ^11,12^. However, changes in fibrillar collagens and tumour stiffness in LMS have not been systematically studied.

Cells adhere to fibrillar collagens through receptors at the plasma membrane including integrins and discoidin domain receptors (DDRs) ^13^. The adhesion of integrins and their stabilisation to form mechano-chemical signalling hubs, known as focal adhesions, are favoured in stiff substrates ^14^. Adhesion to fibrillar collagens and integrin signalling are involved in the activation of the mechanosensors and mechanotransducers YAP and TAZ ^15,16^. In cancer cells, integrin signalling and YAP/TAZ induce proliferation, stemness, chemoresistance, and metastasis ^13,17^. Indeed, YAP is frequently active in LM and STSs and is involved in STS tumorigenesis ^18–20^. Furthermore, in endometrial stromal sarcoma and undifferentiated uterine sarcoma, expression of fibrillar collagens and YAP activation are associated with tumour aggressiveness ^21,22^. Simultaneously, by imposing physical constrictions, dense collagen matrices have an inhibitory effect against tumour growth and invasion ^23^. However, cancer cells often acquire high collagen degradation and mechanical remodelling capabilities by which they are able to colonise neighbouring tissues and form distant metastases ^10^. Matrix metalloproteinases (MMPs) are proteins with collagen degradation and remodelling functions that are central in the progression of some STS types ^24^.

Despite their central role in diverse tumour types including LM and other uterine sarcomas, the impact of fibrillar collagens and their downstream signalling on uterine LMS behaviour remains elusive. In this study, we perform an analysis of the content and morphology of fibrillar collagens in MM, LM, LMS, and other uterine sarcomas. We find that LMS tumours present a characteristically low fibrillar collagen density with an increased number of fibre endpoints. At the gene level, we show that fibrillar collagen-related gene expression is not lower in LMS, instead we observe an increase in the expression of MMPs, particularly MMP14. Using collagen-based *in vitro* 3D cultures, we show that MMP14 activity is necessary for LMS proliferation. In addition, we demonstrate that LMS cell proliferation and migration are less dependent on substrate stiffness than in MM and LM. This reduced sensitivity to stiffness is facilitated by the observed elevated basal YAP activity at low stiffness. This study suggests that MMP14 and YAP are fundamental in LMS proliferation revealing them as potential biomarkers and therapeutic targets for this disease.

## Results

### LMS tumours are characterised by low fibrillar collagen density and high fibre endpoints

To understand the function of fibrillar collagens on uterine leiomyosarcoma (LMS), we explored the differences in fibrillar collagen density and network morphology between normal myometrium (MM), uterine leiomyoma (LM), and LMS tissues. Analysis of picrosirius red-stained tissues with the Fiji macro TWOMBLI ^25^ revealed that LMS tissues present a lower fraction of high density matrix (HDM) regions, indicating that LMS tissues have lower collagen density (**Fig. 1a,b**). Moreover, LMS tissues presented increased fibre endpoints and reduced hyphal growth unit (HGU), a measure of the number of endpoints per unit length (**Fig. 1c**). Comparing the MM at distant or tumour-adjacent (0 – 1 mm from the tumour edge) locations showed reduced fractal dimension and endpoints in tumour-adjacent MM, and a non-significant trend to increased fibre alignment and lacunarity and decreased curvature and branchpoints (**Fig. 1c** and **Supplementary Fig. 1**). These alterations in the density and morphology of tumour-adjacent collagen may result from tensile solid stresses originated by the constant increase in tumour volume ^23^.

**Figure 1.**
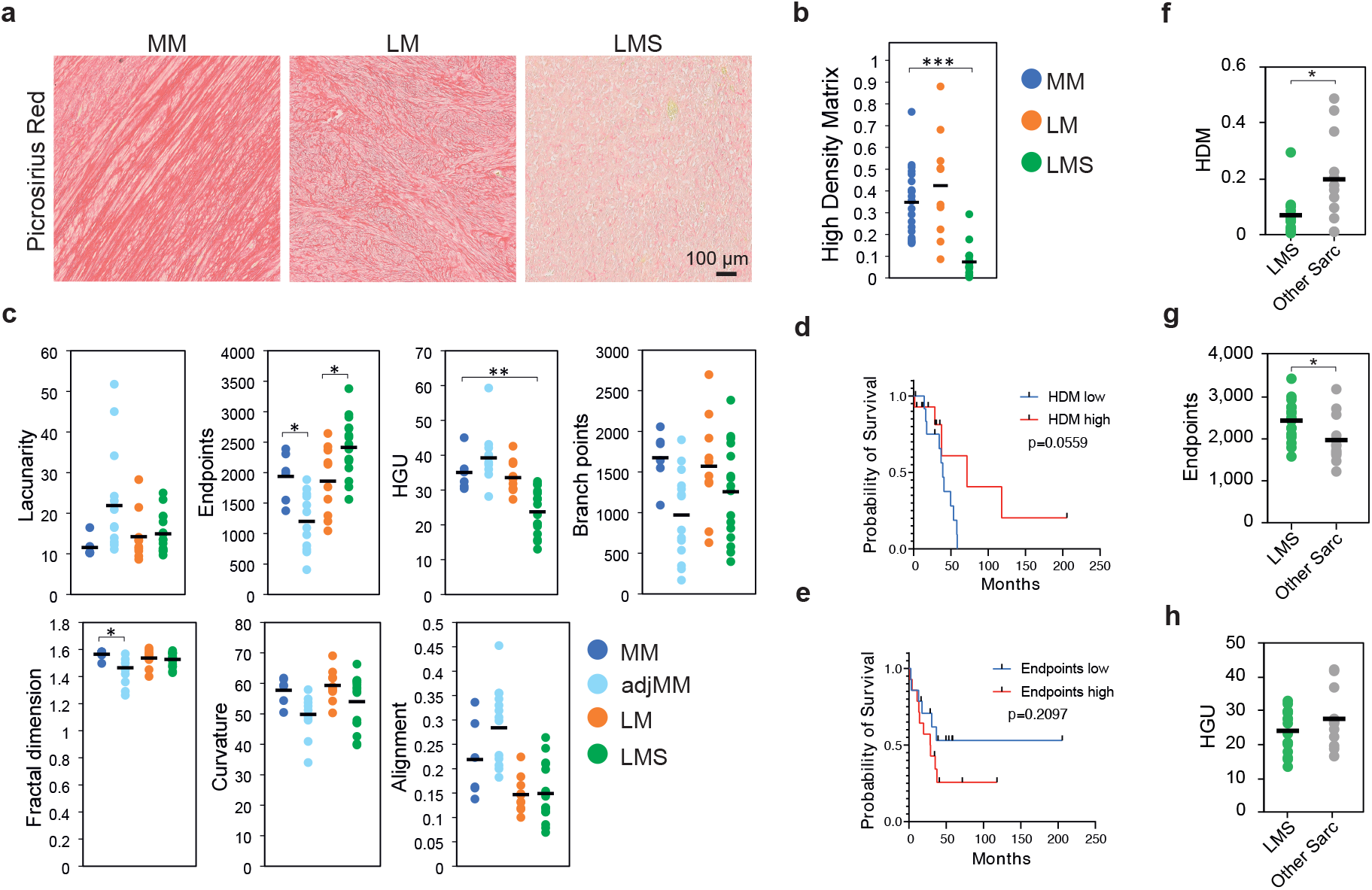
LMS are characterised by low fibrillar collagen density and high fibre endpoints. **a**, Representative images of picrosirius red staining of normal myometrium (MM; n = 6), uterine leiomyoma (LM; n = 10), and uterine leiomyosarcoma (LMS; n = 16) tissues. Scale bar indicates 100 µm. **b**, Quantification of the fraction of high density matrix (HDM) in tissues from (**a**). **c**, Quantification of morphological aspects of the collagenous matrix based on staining from (**a**) and including tumour-adjacent MM (adjMM, <1mm from tumour edge; n = 14). HGU (hyphal growth unit). **d, e**, Kaplan-Meier curves of uterine sarcoma patients grouped based on their fraction of HDM (**d**) or number of fibre endpoints (**e**) (low: n = 14; high: n = 14). **f** - **h**, Quantification of HDM (**f**), fibre endpoints (**g**), and HGU (**h**) comparing LMS tumours (n = 16) and the other sarcomas included in our cohort (other Sarc; n = 11). Each dot in the graphs indicates the average of 4 measurements per tumour sample; horizontal bar indicates the average of the tissue type. * p < 0.05, ** p < 0.01, *** p < 0.001.

To investigate whether the observed differences in fibrillar collagens found in LMS are associated with tumour aggressiveness, we compared the overall survival of a cohort of uterine sarcoma patients (n = 28; **Supplementary Table 1**) presenting distinct HDM and fibre endpoints. Uterine sarcomas with lower HDM and higher endpoints showed a non-significant trend to reduced overall survival (HDM p = 0.0559, endpoints p = 0.2097; **Fig. 1d,e**). Concurrently, the frequency of metastasis was higher in LMS patients with low HDM primary tumours (87.5% in low HDM vs 37.5% in high HDM) and in patients with high endpoints (50% in low vs 75% in high endpoints) (**Supplementary Fig. 2**). To further investigate whether the fibrillar collagen characteristics are a differential trait of LMS, we compared LMS tissues with the other uterine sarcomas included in our cohort. In LMS, HDM was lower and the number of endpoints was higher than in other sarcomas, but no significant differences were observed in HGU (**Fig. 1f-h**). These results indicate that LMS tumours present a particularly low density of fibrillar collagens, and suggest a link between reduced collagen density and tumour aggressiveness.

### MMP14 activity is necessary for LMS cell proliferation in collagenous microenvironments

To explore the origin of the reduced collagen density found in LMS tumours, we compared the expression of fibrillar collagens and collagen biosynthesis and crosslinking-related genes in MM, LM, and LMS tissues. The expression of 3 out of 11 fibrillar collagen and 8 out of 13 collagen biosynthesis genes was upregulated in LMS compared to MM and LM (**Fig. 2a,b**), suggesting that the low density of fibrillar collagens observed in LMS is not caused by a reduced expression of fibrillar collagen-related genes. Next, we investigated whether LMS tumours present differential expression of collagen-degrading MMPs. Compared to MM and LM, LMS tumours showed increased expression of 4 out of 6 genes encoding collagen-degrading MMPs, including *MMP14* as the most highly expressed MMP in LMS (**Fig. 2c**). Similarly, compared to other uterine sarcomas, *MMP14* expression was upregulated in LMS (**Fig. 2d**). These results were confirmed at the protein level, showing higher MMP14 expression in LMS than in benign LM tumours (**Fig. 2e,f**).

**Figure 2.**
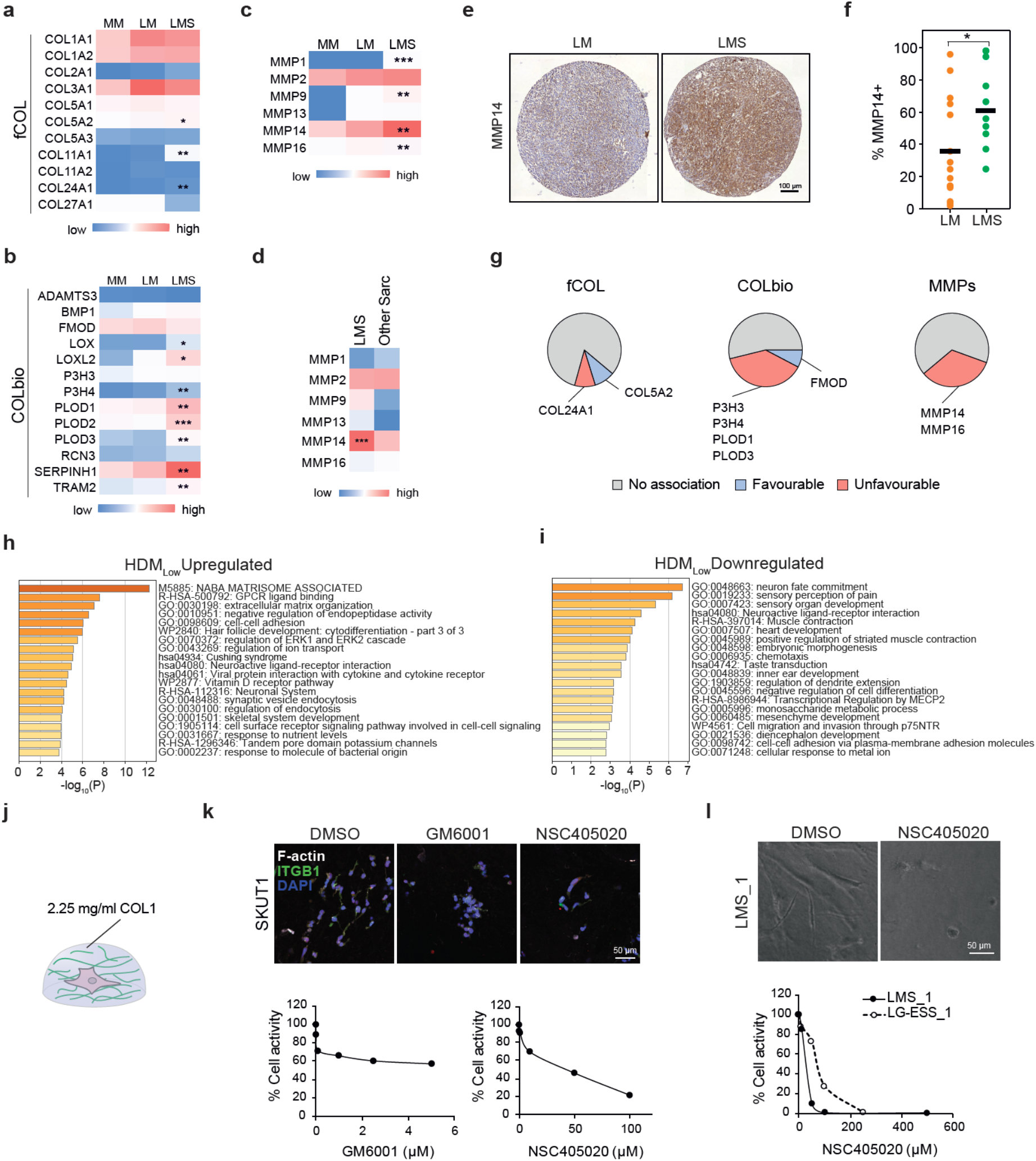
LMS cells highly express MMP14 and depend on its activity for proliferation in collagenous environments. **a** - **c**, Relative average expression of fibrillar collagen (fCOL) (**a**), collagen biosynthesis and crosslinking (COLbio) (**b**), and collagen-cleaving matrix metalloproteinases (MMPs) (**c**), in normal myometrium (MM; n = 31), uterine leiomyoma (LM; n = 28), and uterine leiomyosarcoma (LMS; n = 17). **d**, Relative average expression of MMPs in LMS and other uterine sarcomas included in the cohort (other Sarc; n = 17). **e**, Representative examples of MMP14 protein staining in LM and LMS tissues. Scale bar indicates 100 µm. **f**, Quantification of the percentage of the area of the tissue positive for MMP14 in LM and LMS tissues, indicating higher MMP14 protein expression in LMS. Each datapoint represents one patient; horizontal bars indicate the average per tissue type. **g**, Association between high expression of fCOL, COLbio, and MMP genes and LMS patient prognosis of the TCGA cohort (n = 24). Patients groups were generated by the best cutoff method. **h, i**, Metascape pathway analysis of the genes upregulated (**h**) and downregulated (**i**) in LMS tumours with low fraction of high density matrix (HDMl^ow^; n = 10) compared to HDM^high^ (n = 8) LMS tumours. **j**, Schematic representation of the 3D collagen-based models used to embed cells in (**k**) and (**l**). **k**, Representative images (top) and quantification (bottom) of the response of SKUT1 cells to pan-MMP inhibition (GM6001) or the specific inhibition of MMP14 (NSC405020) based on total ATP content. Each data point indicates the average of 3 technical replicates. Scale bar indicates 50 µm. **l**, Representative images of patient-derived LMS cells (scale bar indicates 50 µm) and quantification of the response of LMS and low-grade endometrial stromal sarcoma (LG-ESS) cells to NSC405020. Each data point indicated the average of 3 technical replicates. * p < 0.05, ** p < 0.01, *** p < 0.001.

Next, we evaluated the prognostic value of fibrillar collagens, collagen biosynthesis-related and MMP gene expression in LMS. High expression of 2/11 fibrillar collagen genes was associated with patient survival (*COL24A1* unfavourable and *COL5A2* favourable association), whilst 4/13 collagen biosynthesis/crosslinking genes were associated with unfavourable prognosis (*P3H3, P3H4, PLOD1, PLOD3*) and 1/13 with favourable prognosis (*FMOD*) (**Fig. 2g**). The expression of *MMP14* and *MMP16* was also associated with unfavourable prognosis in LMS patients. Thus, the expression of various collagen-remodelling and degrading genes is linked to LMS aggressiveness. To further understand the origin of reduced fibrillar collagen density in LMS, we compared the gene expression of LMS tumours with high and low HDM. Pathway analysis of genes upregulated in low HDM tumours indicated that the “NABA Matrisome Associated” was the most significantly altered gene set followed by “GPCR ligand binding”, and “extracellular matrix organisation” (**Fig. 2h**). Amongst the upregulated matrisome-associated genes in low HDM LMS tumours, we found the ECM-degrading MMP13 and ADAMTS20 (**Supplementary Table 2**), further suggesting increased ECM proteolysis in LMS tumours. Contrarily, pathway analysis of genes downregulated in low HDM LMS tumours indicated “neuron fate commitment”, “sensory perception of pain”, and “sensory organ development” as the most significantly altered pathways, and did not reveal any ECM-related pathway (**Fig. 2i**). These results indicate that the low collagen density in LMS tumours is coupled with an enhanced collagen remodelling and degradation.

Finally, to evaluate the function of MMPs and, in particular, MMP14 in LMS, we embedded LMS cells in 3D collagen I-based matrices, mimicking collagen-rich environments such as the MM, followed by MMP pharmacological inhibition (**Fig. 2j**). In the LMS cell line SKUT1, MMP inhibition with the broad-spectrum inhibitor GM6001 affected cell distribution, inducing the formation of cell clusters compared to the single-cell invasive growth pattern observed in the control (**Fig. 2k**). This single-cell growth pattern in SKUT1 cells suggests that these cells disseminate individually within the collagen matrix, reflecting a more aggressive behaviour than cells growing as non-invasive clusters. Specifically inhibiting MMP14 with NSC405020 caused a more pronounced reduction in SKUT1 cell growth, as observed by the lower cell density and reduced total viability, compared to GM6001 (**Fig. 2k**). Next, we evaluated the response of patient-derived LMS cells to MMP14 inhibition and compared it with the response of less aggressive low-grade endometrial stromal sarcoma (LG-ESS) cells. Inhibition of MMP14 in LMS caused a drastic reduction in cell spreading and growth, indicated as reduced total ATP content, an effect that was less pronounced in the LG-ESS cells (**Fig. 2l**). Altogether, these results indicate that, as in various carcinoma and in fibrosarcoma cells, MMP14 activity is essential for the proliferation and aggressive growth pattern of LMS cells in collagen-rich environments ^26^.

### Proliferation of LMS cells is characterised by reduced substrate stiffness-dependence

Cell adhesion to a stiff ECM promotes cell proliferation through integrin signalling and other mechanically-activated pathways ^27^. As LMS tumours present low fibrillar collagen density, we hypothesised that LMS cell proliferation is less dependent on substrate adhesion and substrate stiffness than normal MM and benign LM cells. To investigate the effect of fibrillar collagens on LMS cell proliferation, we first compared the cell proliferation in LMS tumours with normal MM and benign LM. As expected, LMS tumours showed enhanced proliferation index, calculated as the combined expression of multiple proliferation-related genes ^28^, and higher mitotic number than MM and LM tissues (**Fig. 3a,b**). Comparing the proliferation of LMS tumours with high and low HDM showed no differences in proliferation index or mitotic number (**Fig. 3c,d**), indicating that the collagen density of LMS tumours does not correlate with cell proliferation *in vivo*. To functionally evaluate the effect of substrate adhesion and stiffness on LMS cell proliferation, we seeded SKUT1 cells or normal human smooth muscle cells (SMCs) on collagen I-functionalized polyacrylamide hydrogels of increasing stiffness, from very soft 0.5 kPa to supraphysiologically stiff 115 kPa (**Fig. 3e**). The proliferation of SKUT1 cells, indicated by EdU incorporation, showed little variations in the different substrate stiffness (**Fig. 3f,g**). However, SMCs showed a progressive increase in EdU+ cells with increasing substrate stiffness (**Fig. 3f,h**). Similarly, while EdU incorporation steadily increased with substrate stiffness in patient-derived MM cells (n = 4 donors), this effect was less pronounced in LM (n = 3 donors) and LMS cells (n = 4 donors; **Fig. 3i,j**). In general, the highest proliferation was achieved at physiological stiffness (> 4.5 kPa). However, LMS cell proliferation showed high variability between donors, suggesting that the proliferation of some LMS cells is more substrate stiffness-dependent than others. Altogether, these results show that the proliferation of LMS cells presents reduced dependency on collagen adhesion and substrate stiffness.

**Figure 3.**
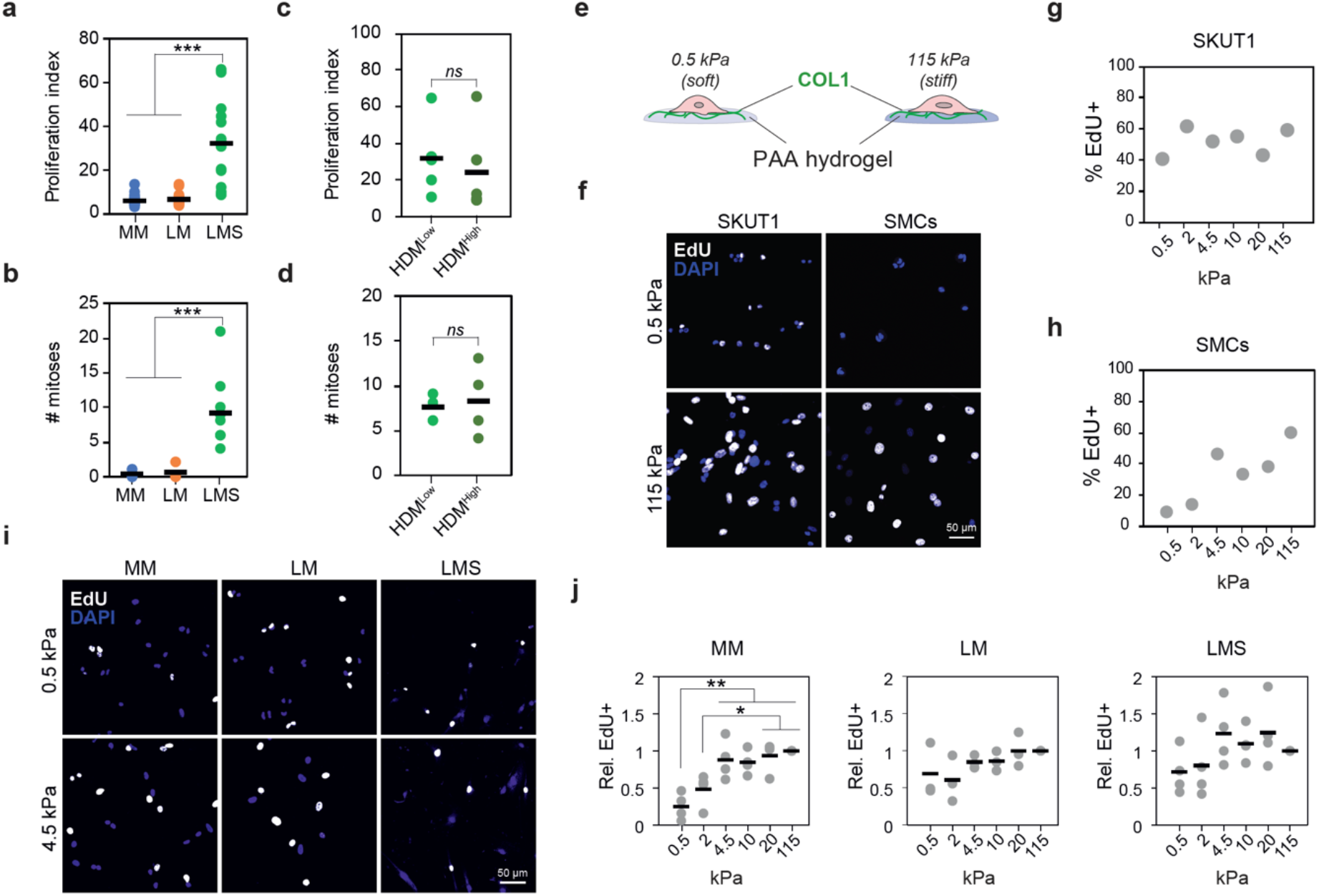
Adhesion to collagen substrates has minimal effect on LMS cell proliferation. **a, b**, Gene expression-based proliferation index (**a**) and number of mitoses per field (**b**) in MM, LM, and LMS tissues indicating enhanced proliferation in LMS. **c, d**, Proliferation index (**c**) and number of mitoses per field (**d**) in LMS tumours with high and low HDM indicating no significant differences between groups. Each dot represents one patient tissue; horizontal bars indicate average per group. **e**, Schematic representation of the 2D model of controlled collagen substrate stiffness based on polyacrylamide (PAA) hydrogels. **f** - **h**, Representative images (scale bar indicates 50 µm) and quantification of cell proliferation measured by EdU incorporation in SKUT1 cells (**g**) and primary smooth muscle cells (SMCs) (**h**) at indicated PAA hydrogel stiffness. **i, j**, Representative images (**i**) and quantification (**j**) of EdU incorporation in MM cells (n = 4 donors), LM (n = 3 donors), and LMS (n = 4 donors) cells adhered to =PAA hydrogel substrates with indicated stiffness. Each dot indicates the average per donor; horizontal bars indicate the average per tissue type. *ns* p > 0.05, * p < 0.05, ** p < 0.01, *** p < 0.001.

### Proliferation at low stiffness is supported by enhanced YAP activity in LMS cells

To further evaluate the lower dependence of LMS cells to collagen adhesion and substrate stiffness, we investigated the subcellular localisation of the mechanotransducers and proliferation regulators YAP/TAZ, found in the nucleus in their active form. Nuclear localisation of YAP/TAZ in SKUT1 increased with stiffness (**Fig. 4a,b**). Similarly, YAP/TAZ nuclear localisation was higher with increasing stiffness in SMCs, although their response to increasing stiffness was more pronounced than in SKUT1 (**Fig. 4c,d**). In MM, LM, and LMS patient-derived cells, nuclear YAP/TAZ progressively increased with higher stiffness (**Fig. 4e,f**). However, unlike in MM and LM cells, the average YAP/TAZ nuclear:cytoplasmic ratio in LMS at 0.5 kPa was >1, indicating mostly nuclear localisation (**Fig. 4e,g**). To investigate whether YAP activity is necessary for LMS cell proliferation, we treated SKUT1 cells with the YAP inhibitor verteporfin. Verteporfin treatment reduced the spreading of SKUT1 cells within 3D collagen matrices as well as the colony area, the number of cells per colony, and the percentage of EdU+ cells (**Fig. 4h,i**). Furthermore, the percentage of apoptotic cells, indicated by cleaved-caspase 3 positivity, was reduced by verteporfin, demonstrating that the lower proliferation was not the result of enhanced apoptosis (**Fig. 4j**). Likewise, verteporfin had a strong negative effect on patient-derived LMS cell growth, whilst this effect was less pronounced in the LG-ESS cells used for comparison (**Fig. 4k**). These results indicate that YAP activity is necessary for LMS cell proliferation. Finally, to evaluate whether YAP is active in LMS tumours despite the low collagen density, we analysed the YAP subcellular localisation in LM and LMS tumours. Subcellular localisation of YAP was mostly nuclear in both LM and LMS tumours (**Fig. 4l,m**). Moreover, the expression of YAP target genes in LMS compared to MM and LM tissues did not reveal specific trends, although various differentially expressed genes were observed (**Fig. 4n**). Altogether, these results indicate that the LMS proliferation-supporting YAP activity is less dependent on substrate stiffness than in MM and LM cells. This enhanced basal YAP activity may facilitate LMS growth on soft, low collagen microenvironments.

**Figure 4.**
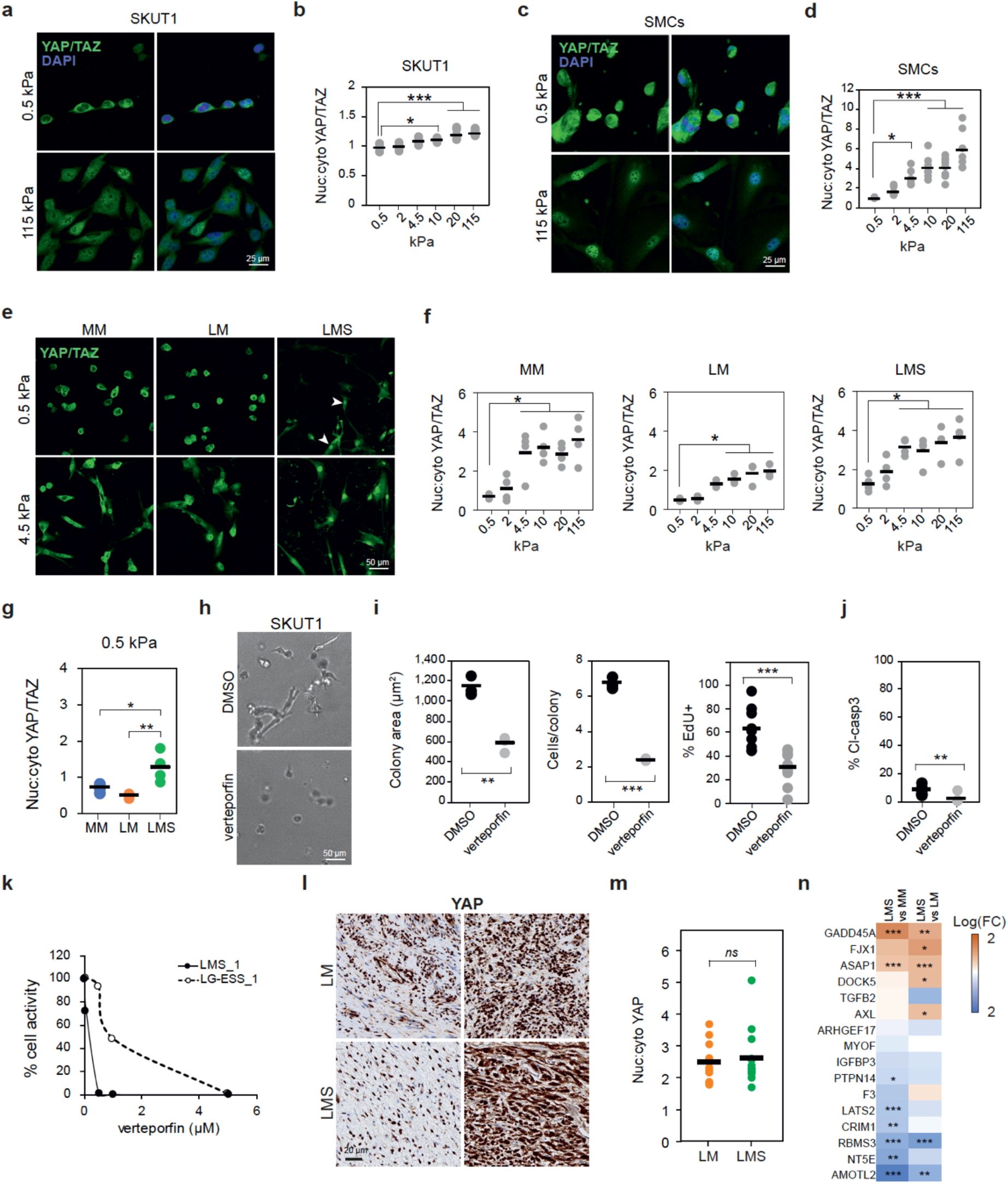
Enhanced YAP activation in LMS supports cell proliferation in soft collagenous substrates. **a, b**, Representative images (**a**) and quantification (**b**) of YAP/TAZ subcellular localisation in SKUT1 cells adhered to PAA hydrogels at indicated stiffness. Scale bar indicates 25 µm. **c, d**, Representative images (**c**) and quantification of YAP/TAZ subcellular localisation in primary smooth muscle cells (SMCs) adhered to PAA hydrogels at indicated stiffness. Scale bar indicates 25 µm. **e, f**, Representative images (**e**) and quantification of YAP/TAZ subcellular localisation in MM, LM, and LMS cells adhered to indicated stiffness. Arrowheads indicate cells with nuclear YAP/TAZ. Scale bar indicates 50 µm. **g**, Comparison of YAP/TAZ nuclear:cytoplasmic ratio in MM (n = 4 donors), LM (n = 3 donors), LMS (n = 4 donors) cells adhered to soft 0.5 kPa PAA hydrogels showing increased nuclear YAP/TAZ in LMS cells. **h**, Representative examples of SKUT1 cells embedded in 3D collagen matrices and treated with YAP inhibitor verteporfin for 48 h. Scale bar indicates 50 µm. **i**, Quantification of colony area, number of cells per colony and EdU incorporation of cells from (**h**) (n = 9). **j**, Quantification of apoptosis indicated by cleaved caspase 3 positivity (cl-casp3) of cells from (**h**). **k**, Quantification of the response of 3D collagen-embedded patient-derived LMS and low-grade endometrial stromal sarcoma (LG-ESS) to verteporfin based on total ATP content. **l, m**, Representative images showing two distinct examples of LM and LMS tissues stained for YAP (**l**) and quantification of nuclear:cytoplasmic YAP ratio showing no difference between the tissues (LM n = 10 and LMS n = 14). Scale bar indicates 20 µm. **n**, Differential gene expression of YAP target genes between LMS and MM and between LMS and LM tissues. * p < 0.05, ** p < 0.01, *** p < 0.001.

### LMS cells present enhanced expression of YAP-activating genes

To investigate whether the enhanced LMS cell YAP activation in low fibrillar collagen microenvironments and on low stiffness substrates derives from enhanced cell-collagen adhesion, we compared the relative cell spreading and adhesion of MM, LM, and LMS cells at distinct stiffness. In all cell types, cell spreading area increased with stiffness, with a significant increase between 0.5 kPa and 4.5 kPa, which approximates the stiffness of normal MM tissue (**Fig. 5a,b**). However, at 4.5 kPa, visible and elongated focal adhesions were only observed in LMS cells (**Fig. 5a**), indicating a lower stiffness threshold for the formation of these signalling hubs. To understand this difference, we compared the gene expression of collagen I receptors between tissue types. Expression of *DDR2, ITGA10*, and *ITGB1* was higher in LMS than in MM and LM (**Fig. 5c**). Next, we investigated whether the collagen substrate stiffness-induced activation of integrin β1 was different in MM, LM, and LMS cells. Expression of active integrin β1 increased in MM and LM cells with stiffness, significantly at >4.5 kPa compared to 0.5-2 kPa (**Fig. 5d**). However, the response of LMS cells from distinct tumours was heterogeneous, although a trend of increasingly active integrin β1 was also observed (**Fig. 5d**). Finally, we compared the levels of active integrin β1 at soft 0.5 kPa in the distinct cell types, which showed a non-significant trend to higher active integrin β1 in LMS cells (p = 0.1813; **Fig. 5e**). These results indicate that there is a higher expression of collagen receptors in LMS cells and a lower stiffness threshold for focal adhesion formation, which could be involved in the enhanced activation of YAP. However, other YAP-activating factors such as the enzymes of the mevalonate pathway 3-Hydroxy-3-Methylglutaryl-CoA Reductase and squalene epoxidase ^29^ also appeared upregulated in LMS at the gene level (**Supplementary Fig. 3**), suggesting that multiple factors contribute to the enhanced YAP activation in LMS.

**Figure 5.**
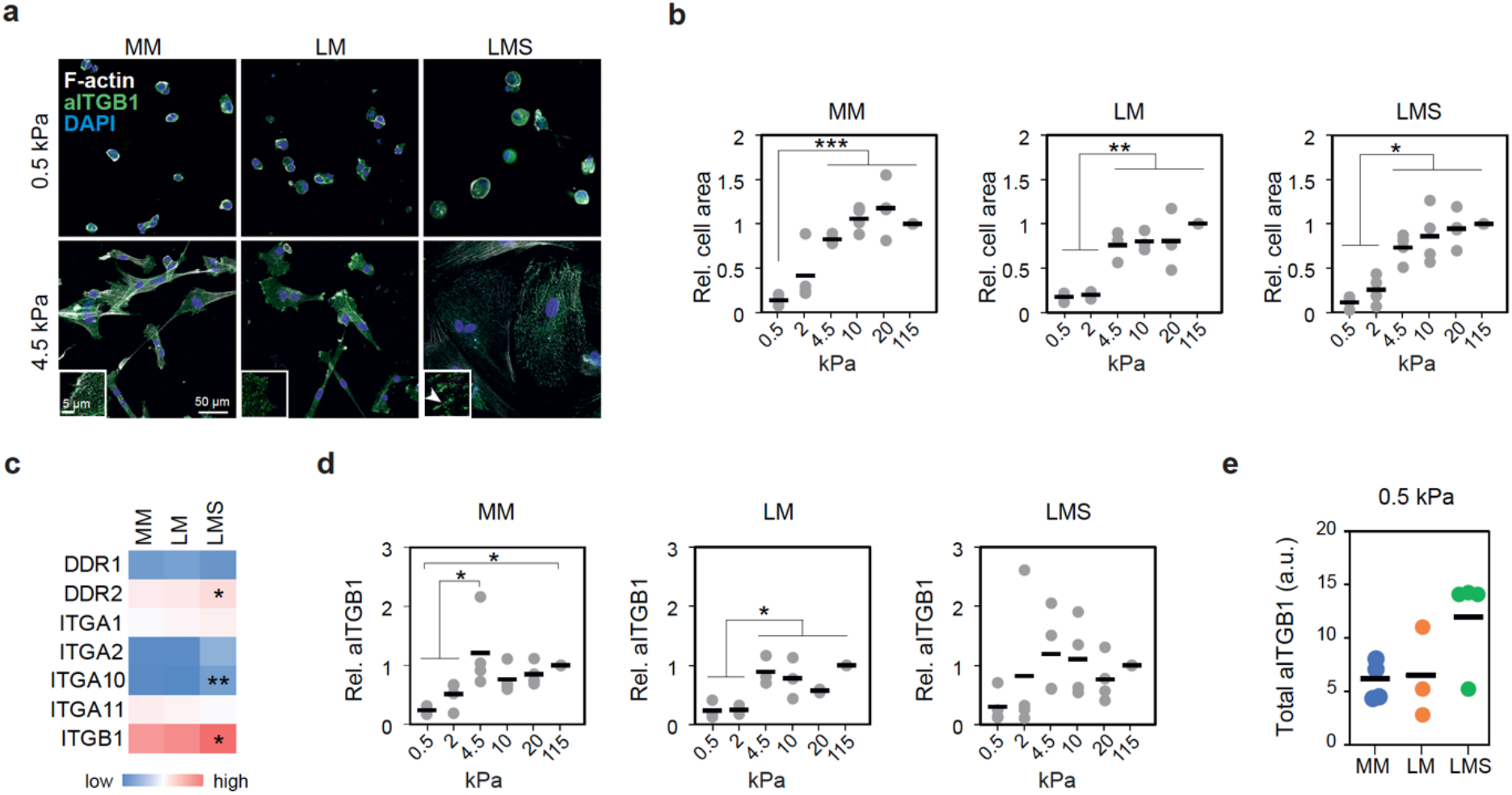
LMS cells overexpress collagen receptors and show enhanced integrin β1 activity. **a**, Representative images of MM, LM, and LMS cells adhered to PAA hydrogels showing distinct morphology and presence of focal adhesions at indicated stiffness. Scale bar indicates 50 µm. Insets show higher magnification of example cellular edges with elongated focal adhesion in LMS (arrowhead). Inset Scale bar indicates 5 µm. **b**, Quantification of spreading cell area of MM, LM, and LMS cells on the indicated substrate stiffness. **c**, Relative collagen receptor gene expression in MM, LM, and LMS tissues showing enhanced expression in LMS of indicated genes. **d**, Change in active integrin β1 intensity (aITGB1) at distinct PAA hydrogel stiffness relative to 115 kPa in MM (n = 4 donors), LM (n = 3 donors), LMS (n = 4 donors) cells. **e**, Comparison of total aITGB1 intensity in MM, LM, and LMS cells at 0.5 kPa showing a non-significant increase in LMS cells. * p < 0.05, ** p < 0.01,

### LMS cell migration shows reduced response to collagen stiffness

Cancer cell migration is a process linked to tumour aggressiveness that is regulated by ECM adhesion and substrate stiffness ^30^. To investigate the effect of substrate stiffness on LMS migration, we seeded patient-derived LMS and LM cells on gels raging from 2 to 20 kPa and tracked them over 13 h. Benign LM cells showed the highest migration speed at 4.5 kPa (**Fig. 6a**). Instead, LMS cell migration speed did not significantly vary across the range of stiffness investigated, although a trend towards reduced speed with increasing stiffness was observed (**Fig. 6b**). Analysis of directionality ratio, a measure that indicates how persistent the migration of a cell is, showed lower directionality ratio of LM cells at 2 kPa (**Fig. 6c**). However, the directionality of LMS cells remained unaffected by substrate stiffness (**Fig. 6d**). Furthermore, the paths followed by LM cells at 4.5 kPa were generally longer than at 20 kPa, whilst this difference was not observed in LMS cells (**Fig. 6e,f**). These results indicate that, within the *in vivo*-relevant stiffness range studied, LM cell migration is stiffness-dependent while LMS migration is stiffness-insensitive.

**Figure 6.**
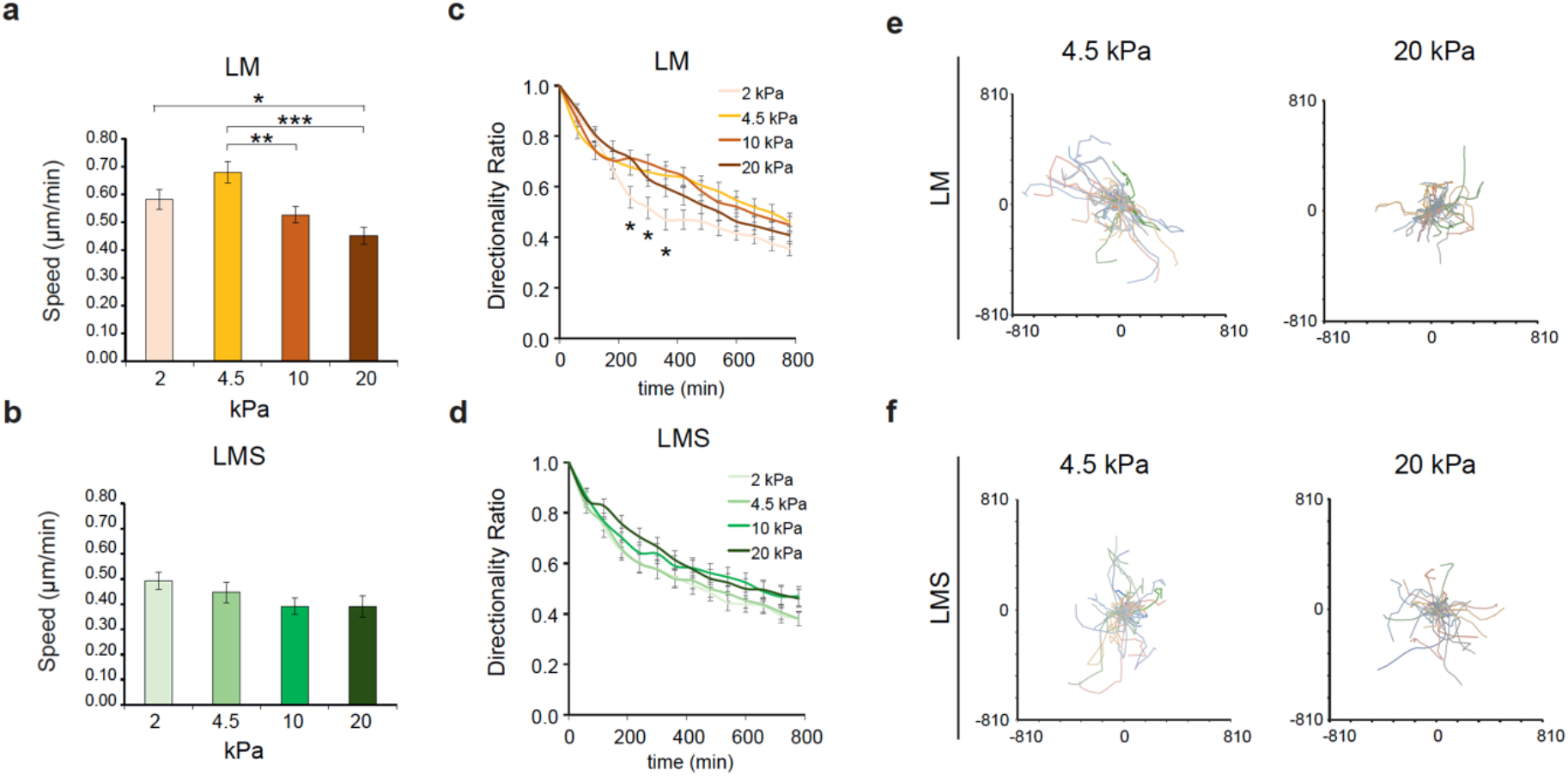
LMS cell migration is not affected by collagen substrate stiffness. **a, b**, Cell migration speed of LM cells (**a**) and LMS cells (**b**) at indicated PAA hydrogel stiffness (n > 10 cells/stiffness per donor; LM n = 3 donors and LMS n =4). **c, d**, Directionality ratio of LM (**c**) and LMS (**d**) cells at distinct PAA hydrogel stiffness. **e, f**, Individual cell trajectories of LM (**e**) and LMS (**f**) cells at indicated stiffness showing relatively shorter trajectories of LM cells at 20 kPa. Bar graphs represent average ±SD. * p < 0.05, ** p < 0.01, *** p < 0.001. *** p < 0.001.

## Discussion

Various clinical trials are currently evaluating the therapeutic potential of targeting collagens and their downstream signalling to treat carcinomas ^31^. These treatments could be an option for LM and specific uterine sarcoma subtypes, as enhanced collagen deposition and YAP activity are associated with undifferentiated uterine sarcoma and endometrial stromal sarcoma aggressiveness ^21,22^. However, due to the lack of knowledge on the relationship between the collagenous ECM and uterine LMS cells, the potential benefit of targeting collagen-related molecules in uterine LMS has not been explored.

Here, we show that, unlike LM tumours, LMS present low fibrillar collagen density with higher fibre endpoints. Moreover, lower collagen density correlates with a trend to reduced overall patient survival and increased probability of metastasis. These results suggest that, unlike the aforementioned carcinomas and non-LMS uterine sarcomas, lower fibrillar collagen expression is associated with tumour aggressiveness. Thus, in principle, LMS patients might not benefit from collagen-reducing therapies. This characteristic of LMS is uncommon but not unique. For instance, in pancreatic cancer, collagen type I deposition is associated with reduced aggressiveness as it acts as a shield against pro-tumour immunity ^32^.

We find that fibrillar collagen-related genes are not downregulated in LMS. Furthermore, we show that LMS tumours highly express various collagen-remodelling genes, including collagen-crosslinking and matrix metalloproteinase genes, particularly *MMP14*. This high expression of collagen biosynthesis and degradation genes suggests that LMS tumours present increased collagen turnover. The enhanced aggressiveness in low collagen-LMS may be given by the high collagen remodelling and degradation, resulting in LMS cell invasiveness, colonisation of neighbouring tissues, and the formation of metastases ^24^. Alternatively, the altered collagenous matrix can lead to increased aggressiveness through the regulation of tumour immunity and other factors of the tumour microenvironment such as the vasculature ^33^, which should be further explored to fully understand the function of collagen and the potential of targeting collagen-related proteins in LMS.

We show that MMP14 inhibition greatly reduces LMS cell proliferation in collagenous matrices and that its gene expression is associated with unfavourable prognosis. The function of MMP14 in LMS proliferation can derive from the reduction of proliferation-inhibiting mechanical confinement as a result of collagen degradation ^34^. Instead, MMP14 activity could be mediating the activation of proliferation-inducing factors such as the heparin-binding EGF-like growth factor, involved in LMS cell survival ^35–37^. Thus, our results indicate that MMP14 is a potential target for the treatment of uterine LMS. However, to date, the usefulness of specific MMP14 inhibitors in cancer treatment has not been demonstrated clinically, and the failure of previous clinical trials has been partly attributed to the inhibition of anti-tumour functions of MMP14 ^31,38,39^.

Enhanced collagen degradation can lead to reduced matrix stiffness and adhesion ligand availability, which are associated with reduced cell proliferation and migration ^27,29,30^. However, here we show that the impact of collagen adhesion on LMS cell proliferation and migration is reduced compared to LM and MM. Interestingly, we show that functional YAP, a central regulator of cell proliferation downstream of ECM adhesion ^13,17^, is necessary for LMS proliferation. Moreover, we find that YAP is overactivated in LMS cells adhered to soft substrates compared to MM and LM. This enhanced YAP activation could derive from the enhanced expression of collagen receptors, the increased activity of other positive regulators of YAP such as the mevalonate pathway, or the reduced activity of YAP-inhibiting molecules ^29,40^. Altogether, these results suggest that inhibiting YAP, either directly or indirectly through YAP-regulatory molecules, could be beneficial for LMS treatment. Moreover, as YAP is involved in chemotherapy resistance, YAP inhibition combined with otherwise poorly effective chemotherapeutics, should be considered ^41,42^.

In conclusion, the results presented here indicate that uterine LMS tumours present a particularly reduced fibrillar collagen content, likely derived from high collagen matrix turnover, associated with tumour aggressiveness. Furthermore, the enhanced YAP activity in LMS tumours supports cell proliferation in their native low-collagen microenvironment. Thus, these results highlight interesting avenues for targeting collagen remodelling and YAP signalling for the treatment of uterine LMS.

## Materials and Methods

### Patient tissue collection (ethical permit)

The retrospective cohort used in this study contains biobanked material with approval (BBK1615). Ethical approval was obtained from the relevant authorities. All cases were reviewed centrally. Patients gave their written informed consent. This study was approved by Stockholm’s regional ethical review board (Dnr2010/1916-31/1).

### Picrosirius red staining

Tissues were deparaffinized and rehydrated through graded ethanol series. Slides were incubated in picrosirius red (PSR) staining solution 1% (w/v) Direct Red 80 (2610-10-8; Sigma-Aldrich) in saturated picric acid (P6744-1GA; Sigma-Aldrich) for 1 h at RT. Slides were washed twice with 0.5% acetic acid, dehydrated in graded ethanol series, cleared in Tissue Clear, and mounted with Pertex (Histolab, Cat # 00811). Slides were imaged with the Zeiss LSM800-Airy confocal microscope using the 561 nm laser. Image quantification was performed with the macro TWOMBLI of Fiji-ImageJ ^25^ and the ridge detection plugin.

### RNA extraction

Fresh frozen tissue samples were collected for RNA extraction. Tissue scrolls were sectioned from OCT-embedded fresh-frozen samples into Buffer RLT with β-mercaptoethanol (Qiagen Sciences LLC, Germantown, MD). RNA was purified following the protocol from RNeasy Micro Kit (Qiagen). RNA concentrations were determined using Qubit RNA HS kit (Thermo Fisher). RNA integrity numbers (RIN) were calculated using the RNA 6000 Pico Kit 2100 Bioanalyzer (Agilent).

### Bioinformatic analysis

Pre-processing, mapping and counting RNA-Seq reads: Each fastq file was assessed using FastQC (v0.11.9) followed by adapter removing and trimming bad quality base calling with Trim-galore (v0.6.6) when it was detected in the quality report. The GRCh38.p13 human genome reference and genome annotation were downloaded (https://ftp.ncbi.nlm.nih.gov/refseq/H_sapiens/annotation/annotation_releases/109.20210226/GCF_000001405.39_GRCh38.p13/) [On May 1st, 2021]. Genome reference was indexed with HISAT2 (v2.1.0) using Hierarchical Graph FM index (HGFM). At this step, only mRNA with “protein-coding” biotype in a complete genomic molecule (RefSeq NC_ format) and transcripts tagged as BestRefSeq were considered. Then, reads were aligned with HISAT2 using the parameter rna-strandness of each sample after inferring the experiment using the script infer_experiment.py from RSeQC (v4.0.0). Samtools was used to filter out reads unmapped, supplementary alignments and reads failing in platform/vendor quality checks and reads with mapping quality below 30 (F=2828 and q=30). Counting reads for each gene was performed using htseq-count (0.13.5) with union mode and the strand-specific information corresponding to each sample.

Data filtering and normalization: Genes were filtered out if less than 50% of each group (MM, LM, LMS, other sarcomas) had count reads below 5. Next, we defined constitutive genes such as genes present in the MM group. Then, genes neither present in constitutive genes nor in LMS or LM groups were removed. DESeq2 (R package) was used to perform a differential analysis and to generate ratios between desired comparisons. Finally, in order to present the data as a heatmap, expression values were calculated as Transcripts Per Million (TPM) using scater (R package).

### Immunohistochemistry

The method for immunohistochemistry (IHC) staining of MMP14 was previously described ^21^. Briefly, TMA sections were deparaffinized and rehydrated. Antigen retrieval was performed using 10 mmol/L sodium citrate pH 6. Endogenous peroxidase was quenched with 0.6% H2O2 for 10 min, 2 × 5 min PBS (for ImmPRESS kit), or with 0.03% H2O2 for 10 min, 1 min H2O, and 10 min PBS [for Tyramide Signal Amplification (TSA) kit]. The ImmPRESS method was used for MMP14 staining with anti-MT1-MMP (LEM) antibody (MAB3328, Millipore, 1:100).

Staining of YAP (ab56701, Abcam, 1:1000) was performed on tissue sections retrieved at the accredited clinical laboratory of the Department of Pathology, Karolinska University Hospital, Sweden. Staining was performed in the routine pathology laboratory by using an automated Ventana Benchmark Ultra system (Ventana Medical Systems, Tucson, AZ, USA).

### Cell culture and isolation

Human uterine leiomyosarcoma-derived SKUT1 cells were cultured in Eagle’s Minimum Essential Medium (EMEM, Thermofisher) supplemented with 10% fetal bovine serum (FBS, Sigma), penicillin/streptomycin (Sigma) and 1x glutaMAX (Gibco) at 37°C and 5% CO_2_. Human penile primary smooth muscle cells and patient-derived primary cells isolated from normal myometrium, uterine leiomyoma, and uterine leiomyosarcoma tumours were cultured in primary Smooth Muscle Cell Basal Medium (Sigma) and penicillin/streptomycin, 10% FBS, 0.5 ng/ml EGF (Sigma), 2 ng/ml bFGF (Sigma) and 5 µg/ml insulin (Sigma) with at 37°C and 5% CO2.

Isolation of patient-derived cells was performed from fresh tissue samples. Tissues were cut into 1-2 mm pieces and subsequently incubated in serum-free medium containing collagenase/hyaluronidase (Stem Cell Technologies) at 37 ºC. After tissue digestion, the solutions were washed with DPBS (Gibco) and red blood cells were lysed with Tris-buffered ammonium chloride solution. After 2X washes with PBS and 2X washes with complete culture medium, cells were seeded on plates coated with 50 µm/ml rat tail collagen I (Sigma).

### Collagen-I polyacrylamide hydrogels

Round microscopy cover slides were washed with 70% ethanol and 0.1 M NaOH, covered with 3-aminopropyltrimethoxysilane (3-APTS, Sigma) for 3 min for activation, incubated for 30 min in 0.5% glutaraldehyde (Sigma), and washed in sterile MilliQ water. Polyacrylamide solutions containing acrylamide monomers (Sigma), crosslinker N,N-methylene-bis-acrylamide (Sigma) and PBS in different concentrations were prepared to create different Young’s modulus (0.5, 2, 4.5, 10, 20 and 115 kPa) as previously described ^43^. 5 µl of 10% ammonium persulfate (Sigma) and 0.75 µl N,N,N′,N′-tetramethylethylenediamine (Sigma) were added into 0.5 ml mixtures, and one drop of the mixture was placed on rain repellent-treated microscopy slides, and the activated cover slides were placed on top. After polymerization (3 to 10 min), the cover slides with polyacrylamide gels were washed with PBS. Next, the cover slides were treated with 1 mg/ml N-sulfosuccinimidyl-6-(4′-azido-2′-nitrophenylamino) hexanoate (Sigma) and exposed to ultra-violet (UV) light to allow subsequent collagen-I binding. The cover slides were incubated at room temperature with 10 µg/ml rat-tail collagen-I (Sigma) for 3 hours, washed with PBS and placed in UV light for sterilization. Wells containing cover slides were seeded with SKUT1 cells or primary SMC, LMS, LM, or MM cells.

### Tissue analysis of mitoses

Per each case, mitotic figures were counted in 10 consecutive high-power fields (400x magnification). The sum of mitotic figures defined the “mitotic count” for each case.

### Immunofluorescence and confocal imaging

Cells were fixed directly on the hydrogel-cover slide in 4% paraformaldehyde for 20 min at RT, permeabilized with ice-cold acetone:methanol (1:1) for 45 s for nuclear staining and blocked in 5% bovine serum albumin (BSA, Biowest) for membrane staining or 5% BSA-0.3% Triton-X (Sigma) for nuclear staining, for 30 min at RT. The cells were incubated with primary antibodies diluted in blocking buffer for 2 hours at RT. After washing, the cover slides were incubated in secondary antibodies and phalloidin for 1 h. EdU staining (Click-iT Plus EdU Imaging Kit, Thermo-Fisher) was performed as described by the manufacturer. Cover slides were washed and mounted on microscopy slides with VECTASHIELD Antifade Mounting Medium with DAPI (Vector Laboratories). Images were taken with a Zeiss LSM 800 confocal microscope with Airyscan and were analysed in ImageJ (Fiji).

### Antibodies

For immunofluorescence, primary antibodies used were Ki67 (8D5; #9449; Cell Signaling Technology; 1:800), active integrin b1 (12G10; ab30394; Abcam; 1:400), YAP/TAZ (63.7; sc-101199; Santa Cruz Biotechnology; 1:200), cleaved-caspase 3 (Asp175; #9664 Cell Signaling Technology; 1:800). Secondary andibodies for immunofluorescence were goat anti-rabbit Alexa Fluor Plus 555 (#A32732, ThermoFisher Scientific; 1:1000) and goat anti-mouse Alexa Fluor Plus 499 (#A32731, ThermoFisher Scientific; 1:1000).

### Live migration analysis

All live cell imaging was conducted in a Cytation 5 imaging reader at +37 °C and 5% CO_2_ (BioTekTM CYT5MPV). Images were taken at 2-hour time intervals over 3 days. For analysis, the manual tracking plugin in ImageJ (Fiji) was used. The distinct cell migration parameters were analysed with the software DiPer ^44^.

## Acknowledgements

We thank the Biomedicum Imaging Core at Karolinska Institutet for imaging facilities. We also thank David Lane and Marie Arsenian-Henriksson for the Cytation 5 microscope.

## Author contributions

J.G.-M. and J.W.C conceived the project. J.G.-M., P.H., and O.G, conducted the in vitro experiments and analysed the data. J.G.-M., R.M.F, and G.K., processed human tissue samples and performed gene expression data analyses. V.Z. analysed tissue mitoses. All authors interpreted the data. T.B.P., K.L., and J.W.C., supervised the study. J.G.-M. wrote the manuscript with contributions from all authors.

## Data and materials availability

All data needed to evaluate the conclusions in the paper are present in the paper and/or the Supplementary Materials. Additional data related to this paper may be requested from the authors.

