## Supplementary Figures for "YAP and collagen remodelling support cell proliferation and tumour aggressiveness in uterine leiomyosarcoma"

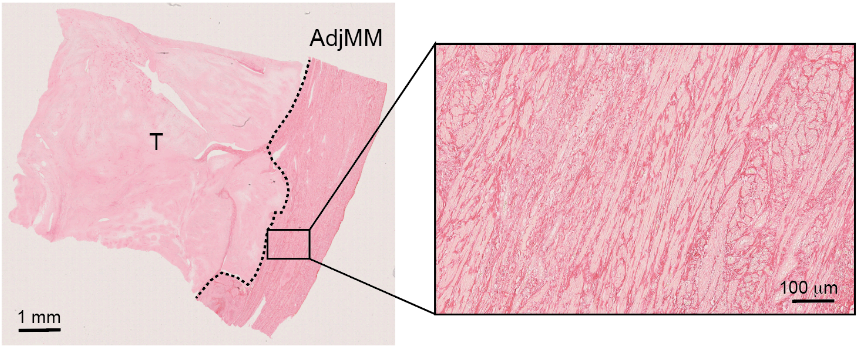


**Supplementary Fig. 1**. Example of tumour tissue (T) with tumour-adjacent myometrium tissue (AdjMM) stained for collagen with picrosirius red showing aligned collagen fibres in AdjMM. Left scale bar indicates 1 mm and the right scale bar indicates 100 μm.

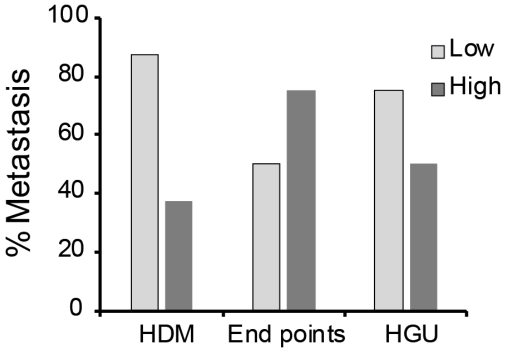


**Supplementary Fig 2**. Percentage of patients presenting metastasis at the time the primary tissue was resected in patients with high (n = 8) and low (n = 8) high-density matrix (HDM), fibre End points, and hyphal growth unit (HGU).

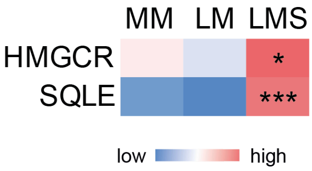


**Supplementary Fig 3**. Expression of the genes of the mevalonate pathway HMGCR and SQLE in MM (n = 31), LM (n = 28), and LMS (n = 17) showing overexpression in LMS. * p > 0.05, *** p > 0.001.
